# Comparison of confound adjustment methods in the construction of gene co-expression networks

**DOI:** 10.1101/2021.05.18.444709

**Authors:** A.C. Cote, H.E. Young, L.M. Huckins

**Affiliations:** Pamela Sklar Division of Psychiatric Genomics, Icahn School of Medicine at Mount Sinai, New York, NY 10029, USA; Department of Psychiatry, Icahn School of Medicine at Mount Sinai, New York, NY 10029, USA; Department of Genetics and Genomics, Icahn School of Medicine at Mount Sinai, New York, NY 10029, USA; Icahn Institute for Genomics and Multiscale Biology, Icahn School of Medicine at Mount Sinai, New York, NY 10029, USA; Seaver Autism Center for Research and Treatment, Icahn School of Medicine at Mount Sinai, New York, NY 10029, USA; Mental Illness Research, Education and Clinical Centers, James J. Peters Department of Veterans Affairs Medical Center, Bronx, NY 10468, USA

**Author notes:** Corresponding author, Correspondence should be addressed to Laura M. Huckins, PhD, Alanna C. Cote, Hannah E. Young.

**Keywords:** Co-expression, Confound, Covariate, Batch effects, RNA-seq, Normalization, Module discovery, Complex traits

## Abstract

Adjustment for confounding sources of expression variation is an important preprocessing step in large gene expression studies, but the effect of confound adjustment on co-expression network analysis has not been well-characterized. Here, we demonstrate that the choice of confound adjustment method can have a considerable effect on the architecture of the resulting co-expression network. We compare standard and alternative confound adjustment methods and provide recommendations for their use in the construction of gene co-expression networks from bulk tissue RNA-seq datasets.

## Background

Large-scale gene expression studies are often subject to technical and biological sources of expression variation including effects of batch, sample characteristics, and environmental factors. Identifying and correcting for confounding sources of expression heterogeneity is a crucial step in data preprocessing and can improve researchers’ ability to quantify biological signal of interest(1,2). Confounding factors can be documented sources of expression variation (known covariates), or derived empirically from the expression dataset (hidden covariates), and adjusting for these factors has become common practice in many population-level gene expression studies. While the benefits of confounding factor correction have been well-characterized in analyses of differential expression and expression quantitative trait locus mapping(2–5), the effects of confounding factor correction on studies of gene co-expression are less well understood (although see (6,7)). This is because confounding factors are difficult to distinguish from gene co-expression, as both variables induce patterns of correlation between genes. In fact, in a well-controlled study, hidden factors are likely to represent biological patterns of gene co-expression in the data(8). Because distinguishing regulatory effects from artifacts is difficult, researchers have historically performed no data correction or known covariate adjustment alone before conducting a co-expression analysis(8–11). More recently, a series of alternative confound adjustment methods have been proposed, designed to correct the expression dataset for confounding factors while retaining patterns of co-expression(7,12,13).

In this study, we evaluate standard and alternative confound adjustment methods in the construction of gene co-expression networks. Using six diverse tissue datasets from the Genotype-Tissue Expression project (GTEx)(3), we identify co-expression networks after adjustment using six data correction approaches(1,7,12,13). To aid researchers in future use of these data correction methods, we present the global and local structure of networks derived from each preprocessed dataset, and assess the accuracy of these networks against two high-confidence human tissue-specific regulatory network references(14,15).

## Results and Discussion

Our analyses were conducted using the subcutaneous adipose, skeletal muscle, spleen, small intestine-terminal ileum, heart-left ventricle, and whole blood tissue datasets from the GTEx project, with sample sizes ranging from 174 to 706 individuals. Each dataset underwent the same preliminary preprocessing including TMM normalization, gene-level filtering, and gene outlier removal. We applied six data correction procedures to each dataset: 1) no correction, 2) known covariate adjustment, 3) probabilistic estimation of expression residuals (PEER)(1), 4) confounding factor estimation through independent component analysis (CONFETI)(12), 5) removal of unwanted variation (RUVCorr)(7), or 6) principal component adjustment (PC)(13). RUVCorr, CONFETI, and PC adjustment are three alternative data correction approaches designed to identify and remove hidden confounds while retaining patterns of coexpression in the dataset. We compare these approaches to one popular standard method of hidden confound adjustment (PEER), known covariate adjustment, and a baseline uncorrected dataset. Description of these methods and their implementation is provided in the Methods section. We generated unsigned weighted co-expression networks for each dataset through calculation of the Pearson correlation between gene pairs, defining an edge as an absolute correlation coefficient > 0.5. We also identified coexpression modules using weighted gene correlation network analysis(16). Module assignments and choice of power parameter for each tissue dataset are provided in Additional Files 2 and 3, respectively.

First, we assessed the impact of each data correction approach on the architecture of resulting coexpression networks through calculation of fundamental network statistics, including node density, clustering coefficients, and standard module properties(17,18). Irrespective of tissue type, CONFETI and PEER adjustment result in the highest proportion of null gene-gene correlations, while known covariate adjustment and no data correction result in the lowest proportion of null gene-gene correlations (Fig. 1A, Additional File 1: Fig. S2). We also find that the degree distribution and clustering coefficients differ between adjustment methods, with genes in CONFETI and PEER-adjusted networks showing fewer network neighbors and higher clustering coefficients as compared to other data correction approaches (Additional File 1: Fig S3-4). Considerable variation in module size, density, and total module number was observed between tissue type and data correction method. Most notably, modules identified from RUVCorr-adjusted data vary widely in size, while modules identified from CONFETI-adjusted data are uniformly-small (Fig. 1C-D). We evaluated the connectedness of genes within co-expression modules through calculation of module density. Similar to module connectivity, module density measures how tight or cohesive genes are within a group, and is equal to the mean adjacency of a module(18). We find that modules identified after PEER adjustment tend to be poorly connected, while modules identified by CONFETI are particularly densely connected (Fig. 1D). Similarity of modules identified after each data correction method is provided in Additional File 1: Figure S5. Overall, there is overlap among modules identified after known covariate adjustment, RUVCorr, PC, and no data correction (Jaccard index>0.5); but not between these and modules identified using CONFETI and PEER.

**Figure 1.**
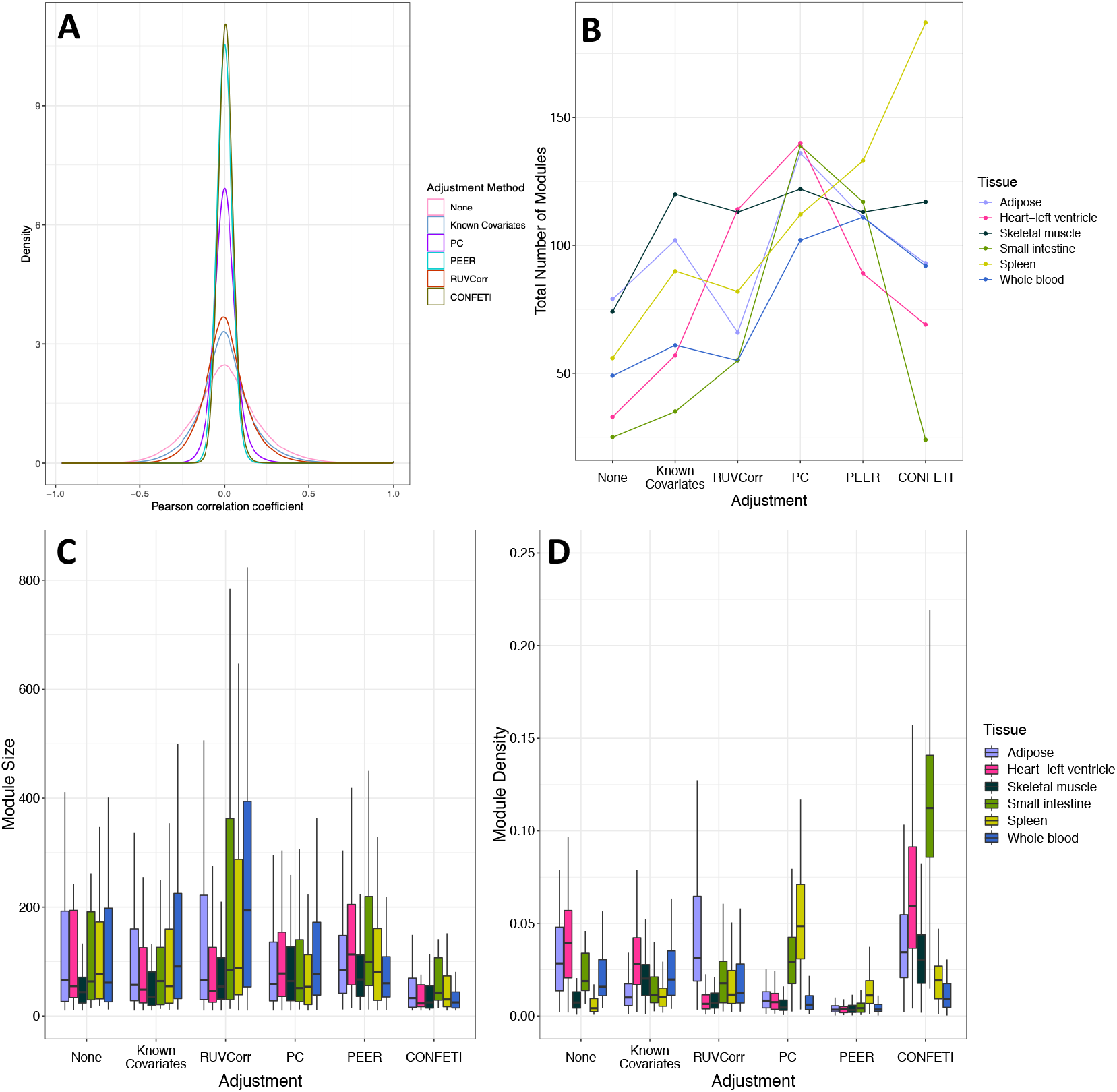
**A)** Distribution of gene-gene correlations for 5,000 randomly-selected genes in the skeletal muscle tissue dataset. **B)** Line plot showing differences in the total number of modules detected across tissue dataset and adjustment method. **C-D)** Box plots showing differences in module size and module density between confound adjustment methods and tissue datasets. Outlier points omitted for ease of visualization.

Next, we evaluated the sensitivity and specificity of each correction method through comparison to two high confidence tissue-specific gene network references. We first employed the framework developed by Somekh et al. (2019)(6), which proposes comparison of gene-gene co-expression to true positive and negative gene pairs obtained from an external network resource. The Genome-Scale Integrated Analysis of Networks in Tissues (GIANT) interface was used as our gene network reference(14). For each expression dataset we: 1) selected high probability true positive and true negative gene pairs from the GIANT tissue network, 2) selected coefficients and FDR-adjusted p-values of Pearson correlation for the corresponding gene pairs in our GTEx expression datasets, and 3) compared adjusted p-values against GIANT network gene pairs to generate receiver operating curves and calculate the area under the curve (AUC). RUVCorr, known covariate adjustment, and no data correction perform similarly on this evaluation metric, while PC, PEER, and CONFETI adjustment result in lower AUC scores than unadjusted data (Fig. 2A).

**Figure 2:**
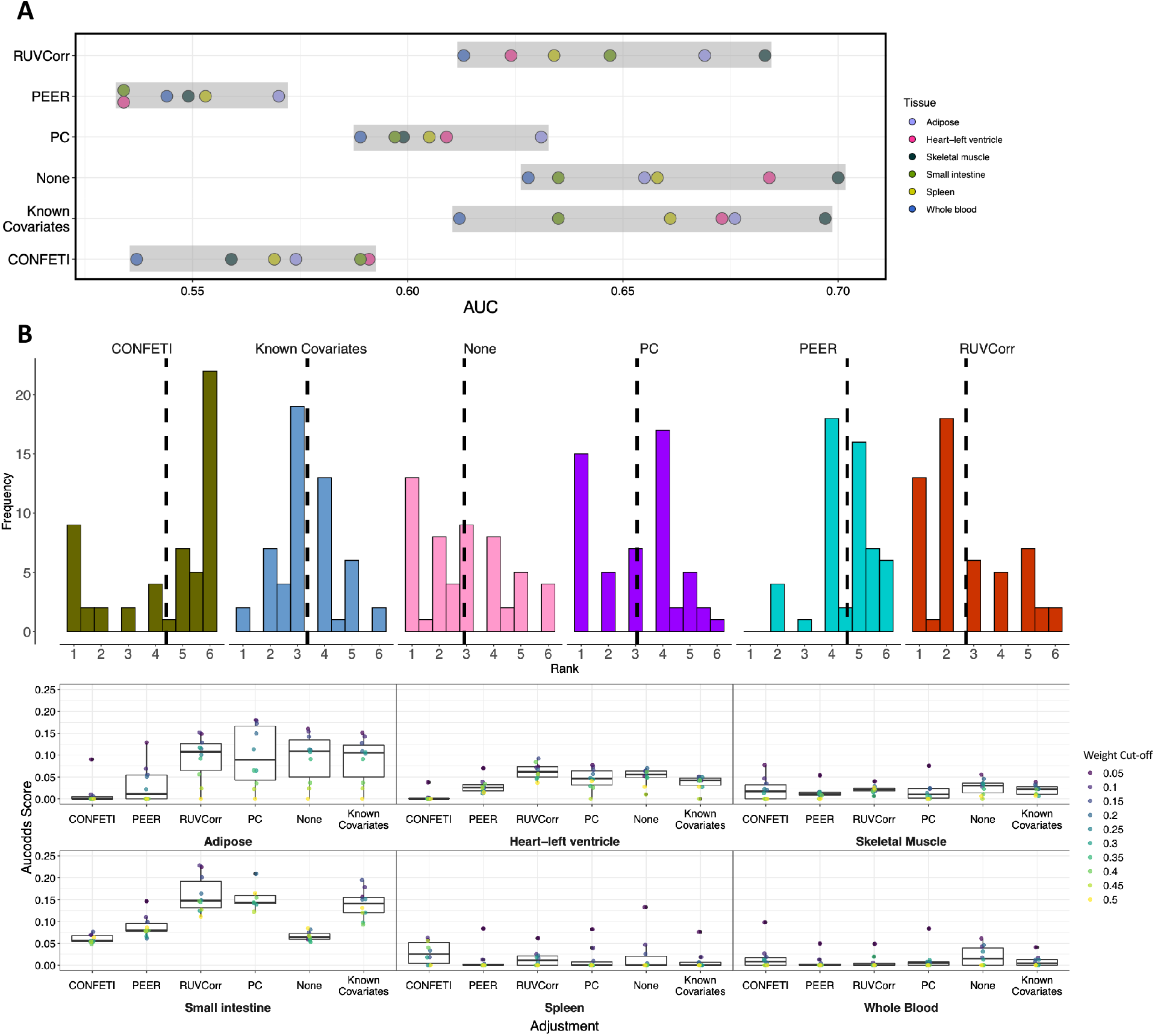
Comparison of covariate adjustment methods. **A)** Area under the curve (AUC) scores for performance evaluation of each adjustment method. Grey boxes highlight the range of scores for each method. **B)** Aucodds scores for performance evaluation of each adjustment method. The top histograms show the distribution of aucodds score rankings by each cut-off-tissue combination (1=best-performing method, highest aucodds score), with dashed lines marking the mean rank. Bottom box plots show the distribution of aucodds scores across tissues and adjustment methods.

Next, we performed comparative network analysis at the modular level, through comparison of coexpression modules in this study and tissue-specific transcriptional regulatory circuits derived from transcription factor motifs, promoter, and enhancer activity information from the FANTOM5 consortium(15,19,20). Co-expression modules were tested for enrichment in gene groups under regulation by the same transcription factor. We summarized the performance of each dataset through calculation of an ‘aucodds’ score(21). This score represents not only the number of target gene groups with a significant enrichment result, but also the extent to which targets of each regulator are enriched in a given module. Although we found substantial variation in performance across tissues, adjustment using RUVCorr, PC, known covariates, and no data correction performed similarly on this evaluation metric, while CONFETI and PEER-adjusted data resulted in poorer overall performance as measured by the average aucodds score ranking (Fig 2B).

Because the CONFETI method was designed to retain only patterns of co-expression associated with common genetic variation, we considered the possibility that CONFETI is effectively capturing a small number of genetically-regulated co-expression modules in our analysis, leading to poor overall representation of the reference gene networks. To investigate this, we calculated the proportion of modules per dataset that showed significant enrichment for targets of at least one regulator (Additional File 1: Fig. S6). Overall, we find that modules identified from CONFETI-adjusted data are less represented in this reference network than other data correction methods.

This study presents multiple lines of evidence suggesting that CONFETI and PEER adjustment may not be appropriate before co-expression network analysis. Both forms of data correction result in particularly sparse networks and weaker representation of two high-confidence gene regulatory networks as compared to other adjustment methods. In addition, networks constructed from PC-adjusted datasets score poorly on one of two reference network comparisons, providing evidence that in some cases PC adjustment may also overcorrect the expression dataset. In this study we adjusted for the number of principal components as suggested by Parsana et al. 2019 in the original proposal of this method. We note however that in some previous co-expression network analyses, adjusting for fewer principal components (typically <10) effectively removes systematic noise from the dataset without overfitting (6,22,23), suggesting that the optimal number of principal components to correct for remains an open question. Lastly, we found that despite differences in network structure, RUVCorr correction, known covariate adjustment, and no data correction all performed similarly in comparison to these two reference regulatory networks. Although we would theoretically expect that correction at least for known technical factors would improve the accuracy of co-expression networks, there is conflicting evidence that this is the case in practice(6,13), and in the present study no form of data correction substantially improved the accuracy of co-expression networks as compared to unadjusted data.

## Conclusions

This study suggests that choice of covariate adjustment can have considerable effects on the structure of the resulting co-expression network. PEER and CONFETI adjustment may overcorrect the expression dataset, removing patterns of biological co-expression of potential interest, and are not recommended for researchers interested in comprehensive co-expression network identification. Conversely, RUVCorr and known covariate adjustment appear to be suitable methods of preprocessing before co-expression analysis, as these methods correct for unwanted effects on expression with no appreciable loss in coexpression signal.

The data correction and module detection methods used in the present study were tested using gene expression from bulk tissue samples. Further research is needed to understand whether these methods are effective in cell-type specific or single-cell expression datasets, as sources of expression heterogeneity and patterns of co-expression likely differ in these data types(24). Further work is also needed to understand whether these results extend to additional methods of network analysis. In particular, alternatives to clustering analysis that capture local co-expression or allow for module overlap warrant evaluation.

## Methods

An illustration of the design of this study is provided in Additional File 1: Fig. S1(25).

### GTEx datasets

We tested performance of each covariate adjustment method using six tissue datasets from the Genotype-Tissue Expression (GTEx) project version 8 release: whole blood, skeletal muscle, spleen, heart-left ventricle, subcutaneous adipose, and small intestine-terminal ileum. Gene read counts and TPMs were downloaded from https://gtexportal.org/home/datasets. Standard RNAseq preprocessing steps were applied to each tissue dataset as follows: 1) TMM normalization of read counts using edgeR and conversion to log2CPM values, 2) gene-level filtering based on threshold of >0.1 TPM in ≥ 20% of samples and ≥ 6 unnormalized reads in ≥ 20% of samples, 3) winsorization of expression values, setting values in specific samples that deviate > 3 standard deviations from other samples to 3 standard deviation limit.

### Covariate adjustment

The following covariate adjustment methods were tested:

1. None: The dataset was not corrected for any known or hidden confounding factor.
2. Known covariates: Information concerning technical factors and sample attributes were downloaded from https://gtexportal.org/home/datasets. Covariates with missing information or zero variance were excluded. We calculated the canonical correlation between remaining covariates; for highly collinear variables, (coefficient > 0.9 for collinear factors) we selected and retained one variable at random. Next, we calculated the variance in expression attributable to remaining continuous technical factors using the *variancePartition* package(26). Lastly, each expression dataset was adjusted for continuous technical factors that explained ≥ 1% variation in ≥ 10% of genes, as well as genotype-derived PC1-5, sex, and binned age. A description of covariates adjusted for in each tissue is provided in Additional File 1: Table S2.
3. Principal components: It has been proposed that for scale-free networks, i.e. networks where the degree distribution follows a power law, patterns of co-expression are sufficiently sparse that principal components of the expression matrix represent an effective form of confound correction(13). As suggested by Parsana et al. (2019), the number of principal components to consider was determined through a permutation-based approach implemented using the “nums.v” function in the *sva* package(13). The number of principal components included in adjustment of each tissue is provided in Additional File 1: Table S1. Significant principal components were regressed on each gene using a linear model and expression residuals obtained.
4. PEER: Probabilistic estimation of expression residuals (PEER) is a popular confound correction method that uses a variant on the traditional factor analysis method to infer hidden factors from the gene expression dataset(1,4). PEER factors were obtained using default settings through the *peer* package. For each dataset we adjusted for the number of PEER factors selected to optimize cis-eGene discovery in the latest quantitative trait locus study by the GTEx Consortium(3): 15 factors for tissues with < 150 samples, 30 factors for tissues with 150-249 samples, 45 factors for tissues with 250-349 samples, and 60 factors for tissues with ≥ 350 samples.
5. CONFETI: Confounding Factor Estimation Through Independent component analysis (CONFETI) is designed to adjust for non-genetic confounding factors while retaining genetically-regulated coexpression (i.e. broad impact eQTL) in the expression dataset(12). Briefly, factors are derived from the full gene expression dataset using independent component analysis. Each independent component is tested for association with genotype in a preliminary broad impact eQTL analysis. Independent components not associated with genotype are considered non-genetic confounding factors and are used in construction of a random effects sample covariance matrix. We identified genetic and non-genetic independent components using the *confeti* package with default settings. Unadjusted gene expression data for each tissue and genotype pruned at R-squared = 0.7 were provided as input. The number of independent components used as confounding factors for each tissue is provided in Additional File 1: Table S1. Five ancestry PCs were regressed on the expression of each gene in a linear mixed model using the *lrgpr* package(27) with non-genetic confounding factors provided as a sample covariance matrix, and gene expression residuals obtained.
6. RUVCorr: The removal of unwanted variation (RUV) method is a multivariate linear model that estimates systematic noise through factor analysis on an expression matrix of empirically-derived negative control genes, i.e., genes in the data with low expression variation that are not expected to be associated with the biological signal of interest (co-expression)(7). In an attempt to mitigate bias in the case where systematic noise and biological signal of interest are correlated, the RUV method uses ridge regression to estimate the effect of systematic noise on expression, and regresses this systematic noise from the expression dataset. The dimensionality of the noise (k) is chosen by the researcher through visual inspection of plots of the distribution of negative and positive control genes in each dataset. A subset of 2,000 genes were used as empirically-derived negative controls while sodium channel genes, major histocompatibility complex genes, and genes that encode for the protein component of the ribosome were used as positive controls (Additional File 1: Fig. S7-S12, positive control gene groups provided in Additional File 4). The ridge parameter (J) is chosen through visual inspection of relative log expression plots (Additional File 1: Fig. S13). Optimal parameters will reduce the correlation between random genes, retain correlation between positive control genes, and best retain gene expression variances in the dataset. RUV correction was applied to each dataset using the *RUVcorr* package, and expression residuals obtained. Choice of RUV parameters for each tissue are provided in Additional File 1: Table S1.

### WGCNA co-expression module search

Co-expression modules were identified using weighted gene correlation network analysis (WGCNA)(16). Each dataset was transformed to a soft-thresholding power ® to approximate scale-free topology (choice of power parameter provided in Additional File 3), followed by construction of an unsigned network with a minimum module size of 10 genes. Modules were merged if correlation of their module eigengenes exceeded a Pearson correlation coefficient of 0.75.

### Comparative network analysis

We used the following methods to compare our co-expression network results to two external gene network references:

1. AUC measure: First, we compared our co-expression network results to high probability true positive and negative gene pairs from the GIANT interface. To obtain high probability gene pairs, for each tissuespecific GIANT network we filtered for genes present in at least one GTEx expression dataset, ranked the network by posterior probability, and kept the top 20,000 and bottom 20,000 gene pairs as true positive and negative gene pairs, respectively. After filtering, 50,405,820 gene pairs remained in each GIANT tissue network, so the selected true positive and negative gene pairs represent the top 0.04% and bottom 0.04% of interactions for each reference network. Not every gene is expressed in each tissue, so fewer gene pairs were tested in the final analysis. The number of true positive and negative gene pairs tested for each tissue are provided in Additional File 1: Table S1. Finally, we calculated the Pearson correlation coefficient and FDR-adjusted p-values for the corresponding gene pairs in the expression dataset, and compared the adjusted p-values against GIANT network gene pairs to generate receiver operating curves and calculate the area under the curve (AUC).
2. Aucodds measure: Next, we tested our identified co-expression modules for enrichment of targets of a shared transcription factor. Using the Marbach et al. (2016) regulatory networks as a reference dataset(15), each gene module was tested for enrichment in target gene groups at various cut-off weights for a true regulator-target gene relationship through a Fisher’s exact test. For each Fisher’s exact test, the background genome size was defined as the number of genes in that particular GTEx expression dataset. For each significant enrichment result (Holm-adjusted p<0.1), we obtained the maximum odds ratio for every regulator across modules. Finally, the performance of each network was summarized through calculation of an ‘aucodds score’(21). The aucodds score is the area under the curve formed by the proportion of regulators with an odds ratio greater than a certain cut-off and the log10 odds ratio cut-off within the OR interval of 1-1000.

## Declarations

Availability of data and materials: Analyses were performed using scripts available at https://github.com/accote45/confound-comparison. Tissue-specific GIANT networks(14) can be downloaded from http://giant.princeton.edu/download/. Marbach et al. (2016) networks constructed from FANTOM5 Consortium data(15) are available as Additional Files 5-10 and can also be downloaded from http://regulatorycircuits.org. The GTEx genome sequencing data used for the analyses in this manuscript were obtained from dbGaP at http://www.ncbi.nlm.nih.gov/gap through accession number phs000424.v8.p2. All other GTEx datasets supporting this article are available in the GTEx repository, at: https://gtexportal.org/home/datasets. The GTEx Project was supported by the Common Fund of the Office of the Director of the National Institutes of Health (commonfund.nih.gov/GTEx). Additional funds were provided by the NCI, NHGRI, NHLBI, NIDA, NIMH, and NINDS. Donors were enrolled at Biospecimen Source Sites funded by NCI\Leidos Biomedical Research, Inc. subcontracts to the National Disease Research Interchange (10XS170), Roswell Park Cancer Institute (10XS171), and Science Care, Inc. (X10S172). The Laboratory, Data Analysis, and Coordinating Center (LDACC) was funded through a contract (HHSN268201000029C) to the The Broad Institute, Inc. Biorepository operations were funded through a Leidos Biomedical Research, Inc. subcontract to Van Andel Research Institute (10ST1035). Additional data repository and project management were provided by Leidos Biomedical Research, Inc.(HHSN261200800001E). The Brain Bank was supported supplements to University of Miami grant DA006227. Statistical Methods development grants were made to the University of Geneva (MH090941 & MH101814), the University of Chicago (MH090951, MH090937, MH101825, & MH101820), the University of North Carolina - Chapel Hill (MH090936), North Carolina State University (MH101819),Harvard University (MH090948), Stanford University (MH101782), Washington University (MH101810), and to the University of Pennsylvania (MH101822).

## Author contributions

AC and LMH conceived the study and designed the experiments. AC, HY, and LMH consulted on code and analytical decisions. AC performed the analysis and wrote the manuscript with input from all co-authors. All authors read and approved the final manuscript.

## Acknowledgements

This work was supported in part through the computational resources and staff expertise provided by Scientific Computing at the Icahn School of Medicine at Mount Sinai. Research reported in this paper was supported by the Office of Research Infrastructure of the National Institutes of Health under award number S10OD018522 and S10OD026880. The content is solely the responsibility of the authors and does not necessarily represent the official views of the National Institutes of Health.

## Supplementary Information

Additional File 1: Supplemental Figures S1-S13. Supplemental Table S1-2. (doc)

Additional File 2: Module assignment for each gene. Each sheet provides module assignments for a particular tissue dataset. (xlsx)

Additional File 3: Choice of power parameter for each co-expression module search. (xlsx)

Additional File 4: Positive control gene groups used for evaluation of RUVCorr parameters. (xlsx)

Additional File 5: Adult blood regulatory circuit(15). (txt)

Additional File 6: Adult heart regulatory circuit(15). (txt)

Additional File 7: Adult skeletal muscle regulatory circuit(15). (txt)

Additional File 8: Adult small intestine regulatory circuit(15). (txt)

Additional File 9: Adult spleen regulatory circuit(15). (txt)

Additional File 10: Adult adipose regulatory circuit(15). (txt)

