## Additional File 1 for "Comparison of confound adjustment methods in the construction of gene co-expression networks"

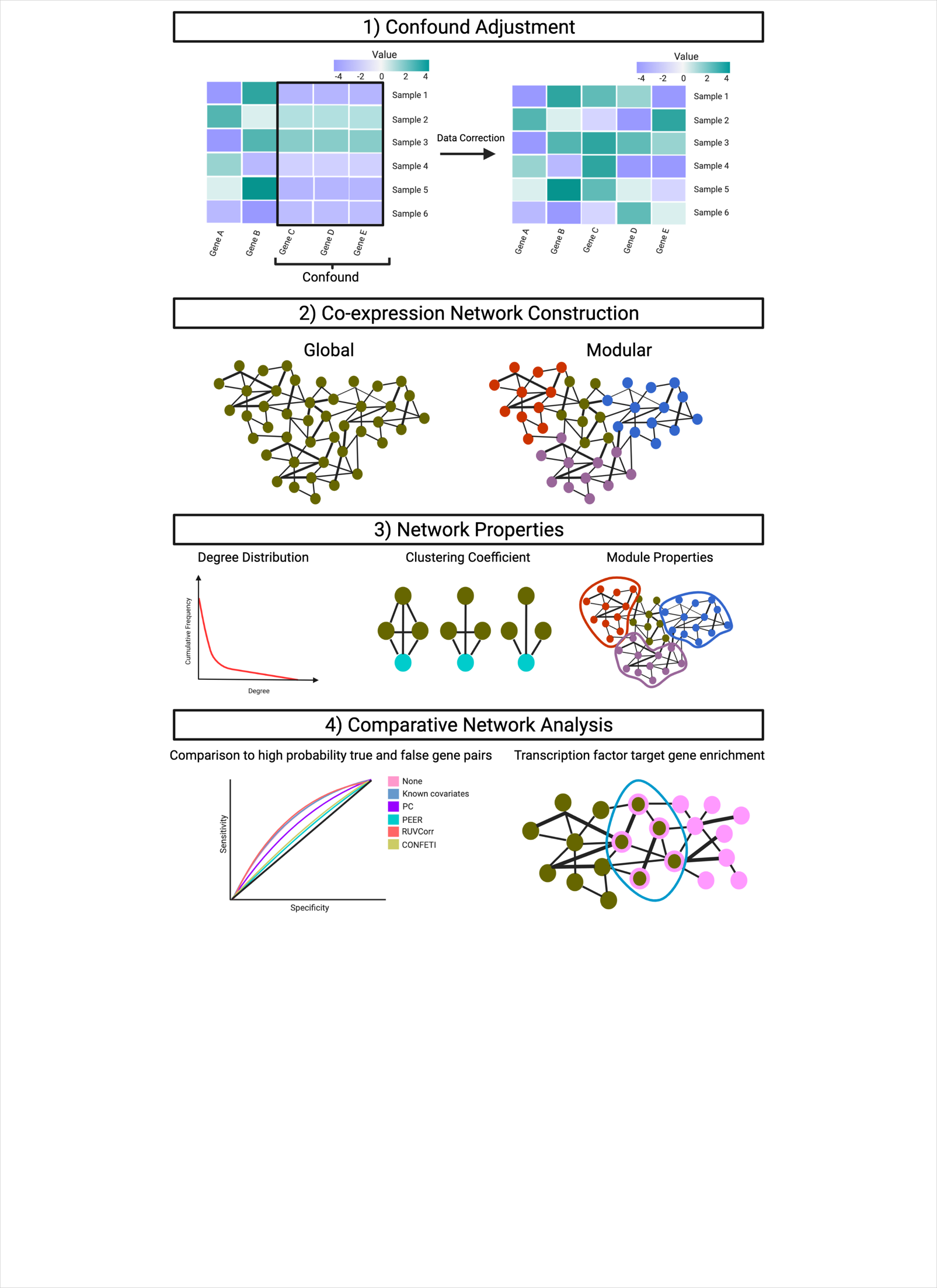

**Figure S1. Illustration of confound adjustment evaluation framework.** Six tissue datasets from the GTEx project were used for this study: whole blood, subcutaneous adipose, spleen, heart-left ventricle, skeletal muscle, and small intestine-terminal ileum. Our framework for assessing confound adjustment was as follows: 1) Adjustment of the gene expression dataset using each data correction approach; 2) Construction of global and modular co-expression networks; 3) Investigation of topological network properties; 4) Evaluation of each data correction approach through comparative network analysis.

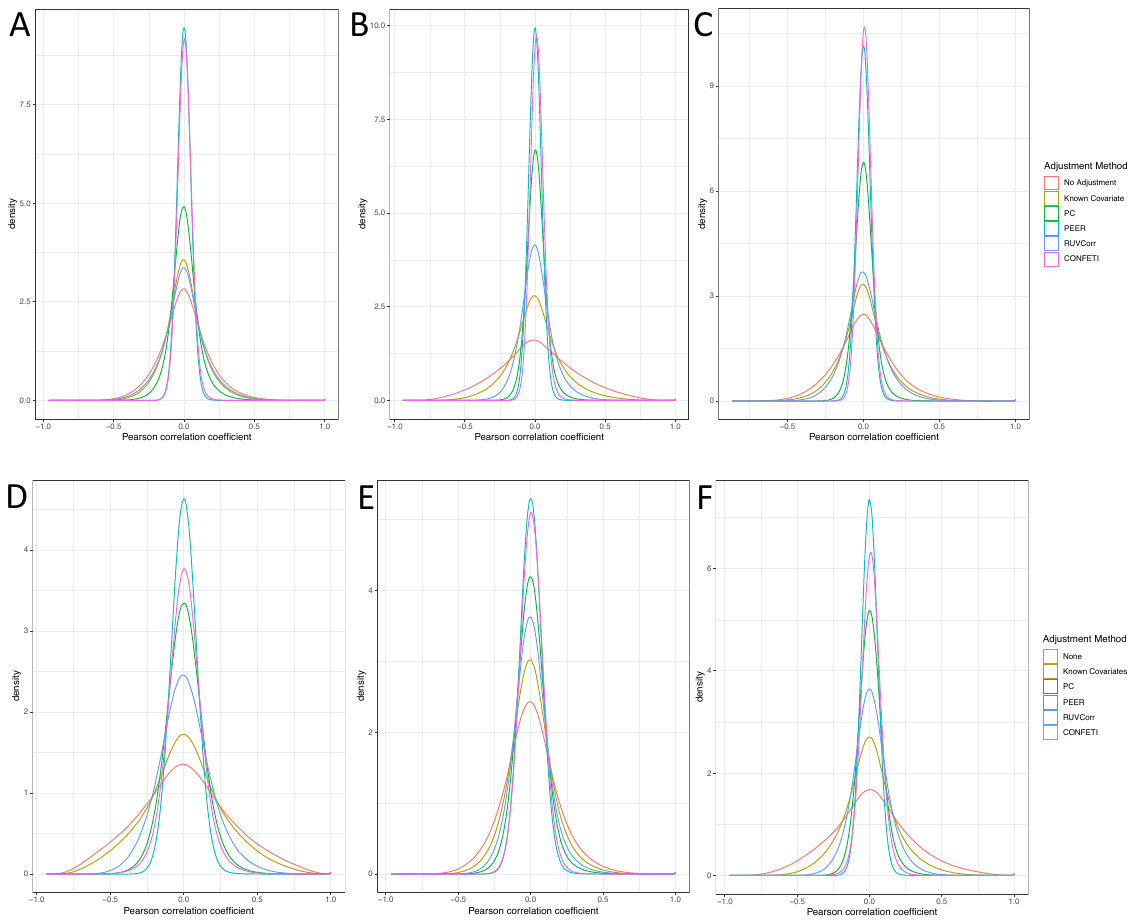

**Figure S2.** Distribution of gene-gene correlation for 5,000 randomly-selected genes. A) Adipose, B) Whole Blood, C) Skeletal muscle, D) Small intestine, E) Spleen, F) Heart-left ventricle

**Figure S3**. Degree distribution of the co-expression network for each data correction-tissue combination. Edges defined as an absolute Pearson correlation coefficient > 0.5. A) Heart-left ventricle, B) Whole blood, C) Spleen, D) Adipose, E) Skeletal muscle, F) Small intestine.

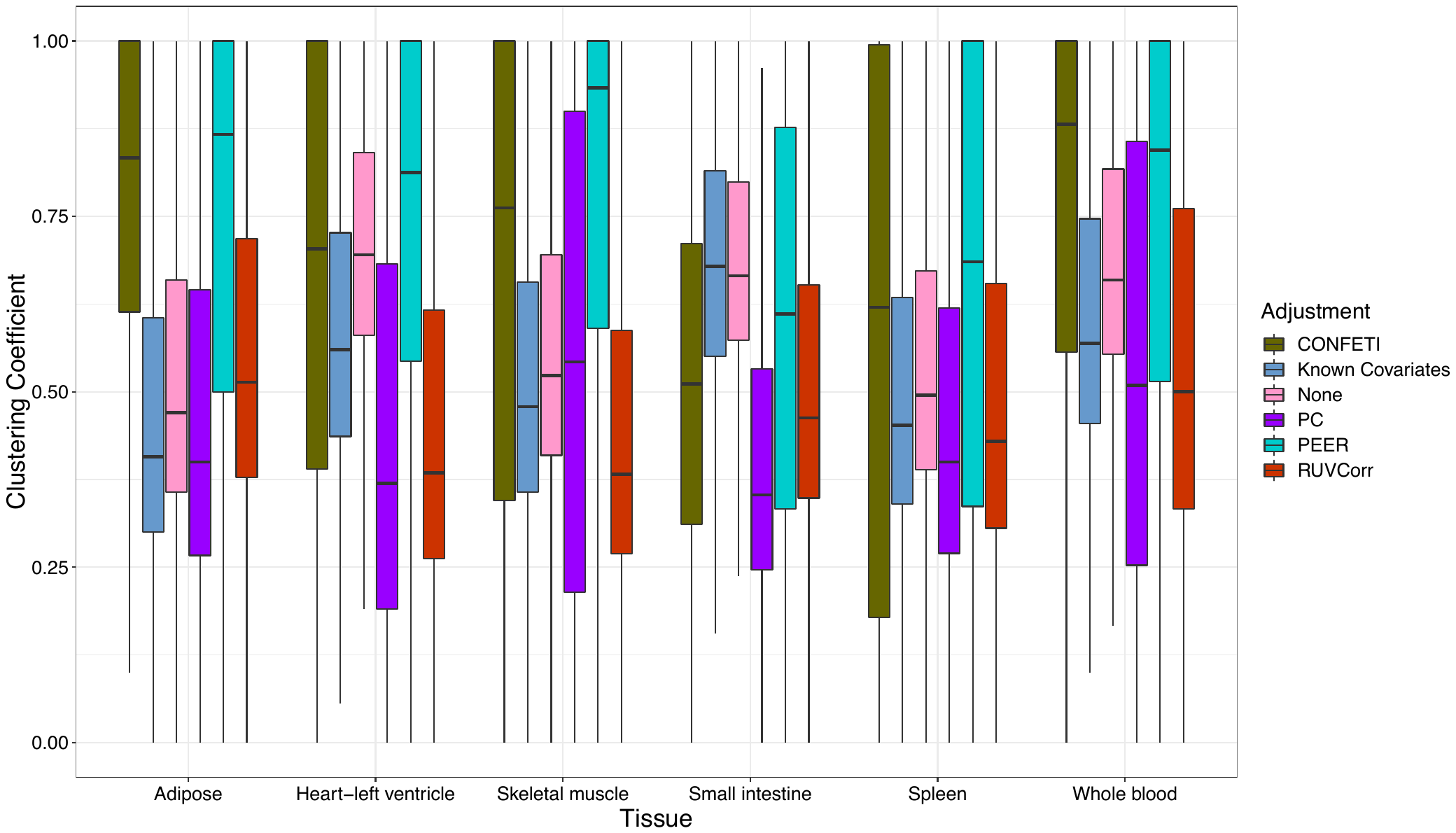

**Figure S4**. Distribution of clustering coefficients for each data correction-tissue combination. Edges defined as an absolute Pearson correlation coefficient > 0.5.

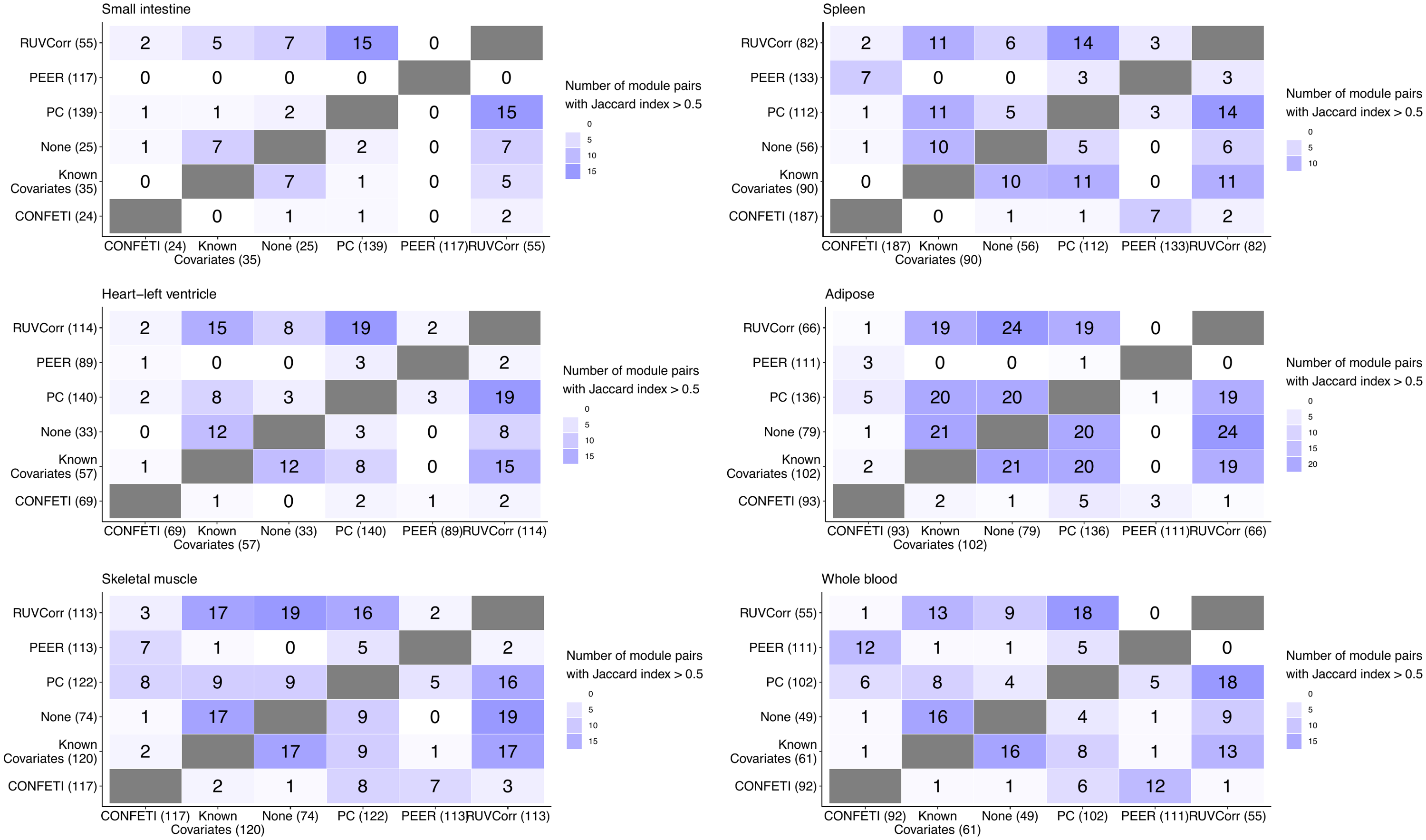

**Figure S5**. Similarity of modules between adjustment methods, as measured by the number of module pairs with Jaccard index > 0.5. Axis label = Adjustment method (Total number of module

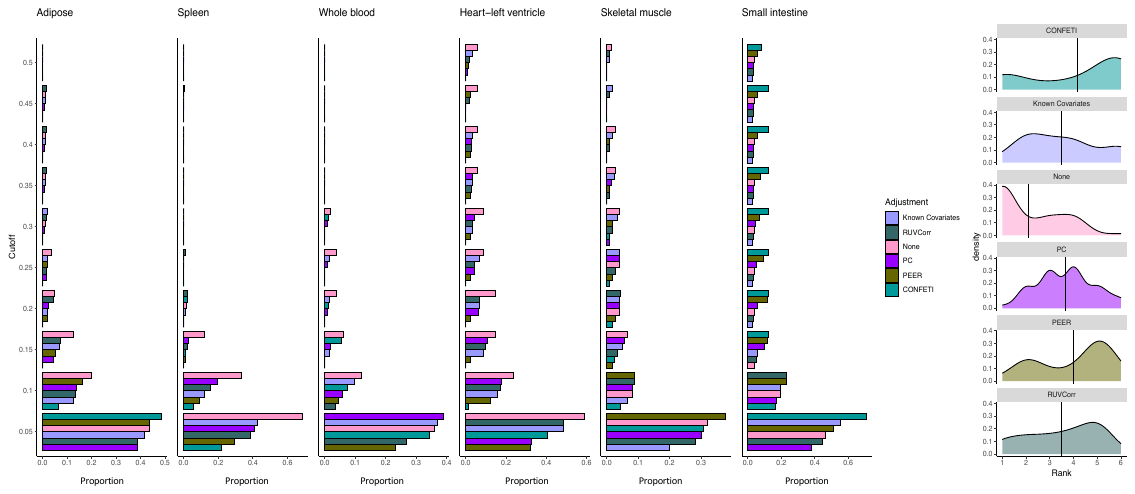

**Figure S6**. Proportion of modules showing significant enrichment (FDR 5%) for the targets of at least one regulator. Y-axis denotes the cut-off weight for a true regulator-target gene relationship. The right density plots show the distribution of proportion rankings by each cut-off-tissue combination (1=best-performing method, highest proportion of modules with a significant enrichment result).

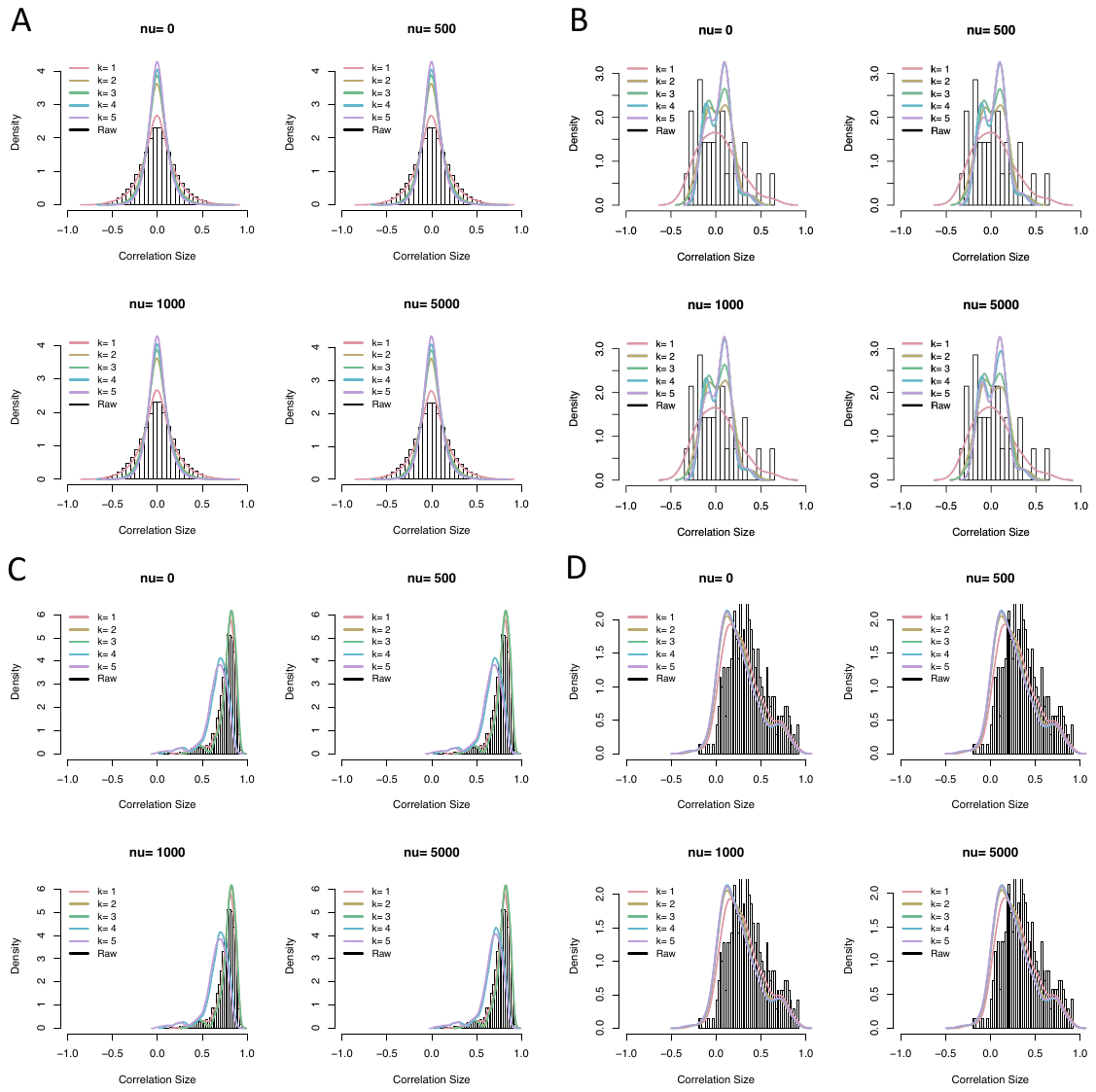

**Figure S7**. Distribution of gene-gene correlations at different RUVCorr parameter settings. Tissue = skeletal muscle. The underlying histogram shows the distribution of gene-gene correlations for each gene set, calculated using unadjusted data. A) Random genes, B) Sodium channel genes, C) Ribosome genes, D) Major histocompatibility complex (MHC) genes

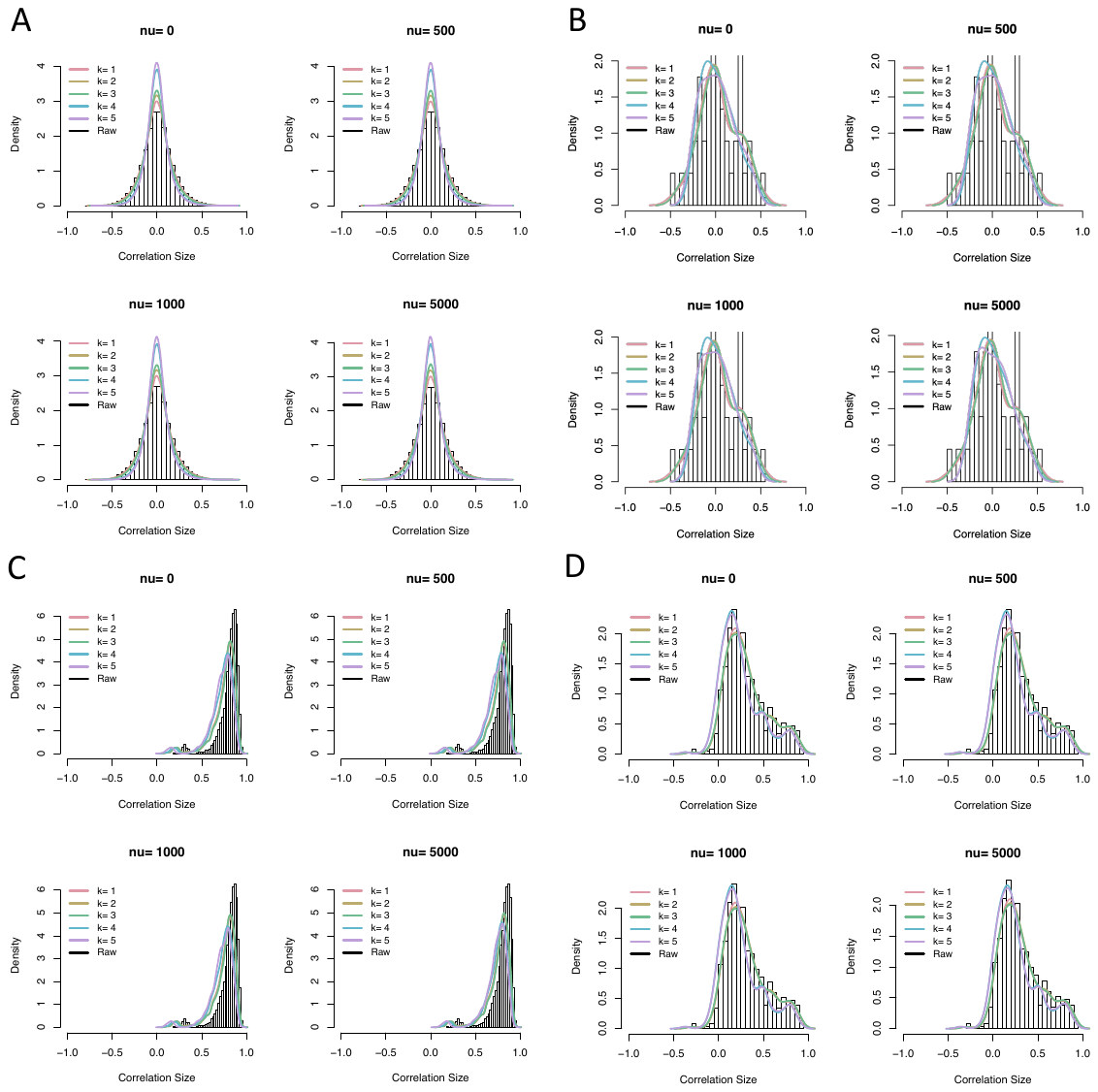

**Figure S8**. Distribution of gene-gene correlations at different RUVCorr parameter settings. Tissue = adipose. The background histogram shows the distribution of gene-gene correlations for each gene set, calculated using the unadjusted data. A) Random genes, B) Sodium channel genes, C) Ribosome genes, D) Major histocompatibility complex (MHC) genes

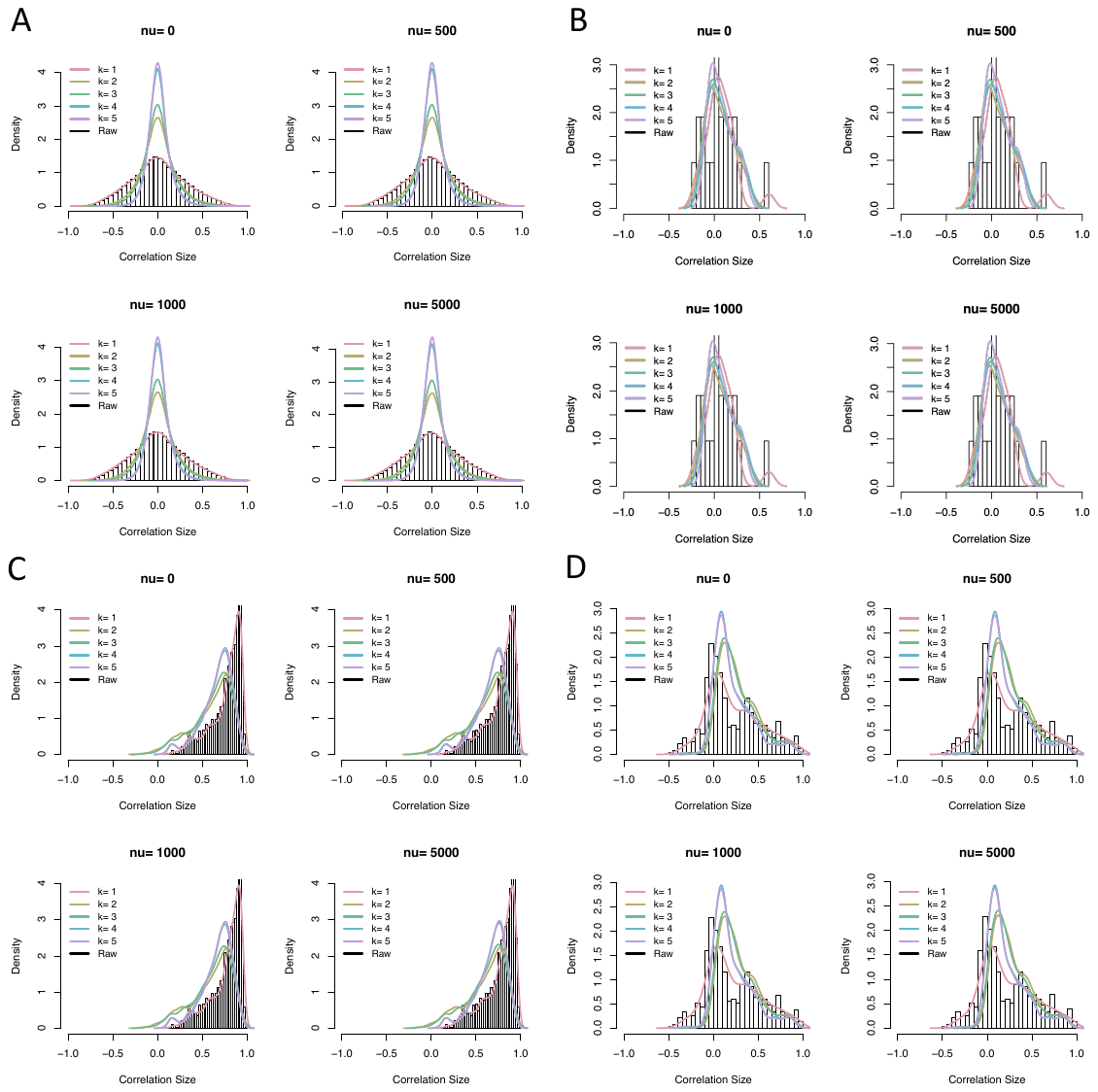

**Figure S9**. Distribution of gene-gene correlations at different RUVCorr parameter settings. Tissue = whole blood. The background histogram shows the distribution of gene-gene correlations for each gene set, calculated using the unadjusted data. A) Random genes, B) Sodium channel genes, C) Ribosome genes, D) Major histocompatibility complex (MHC) genes

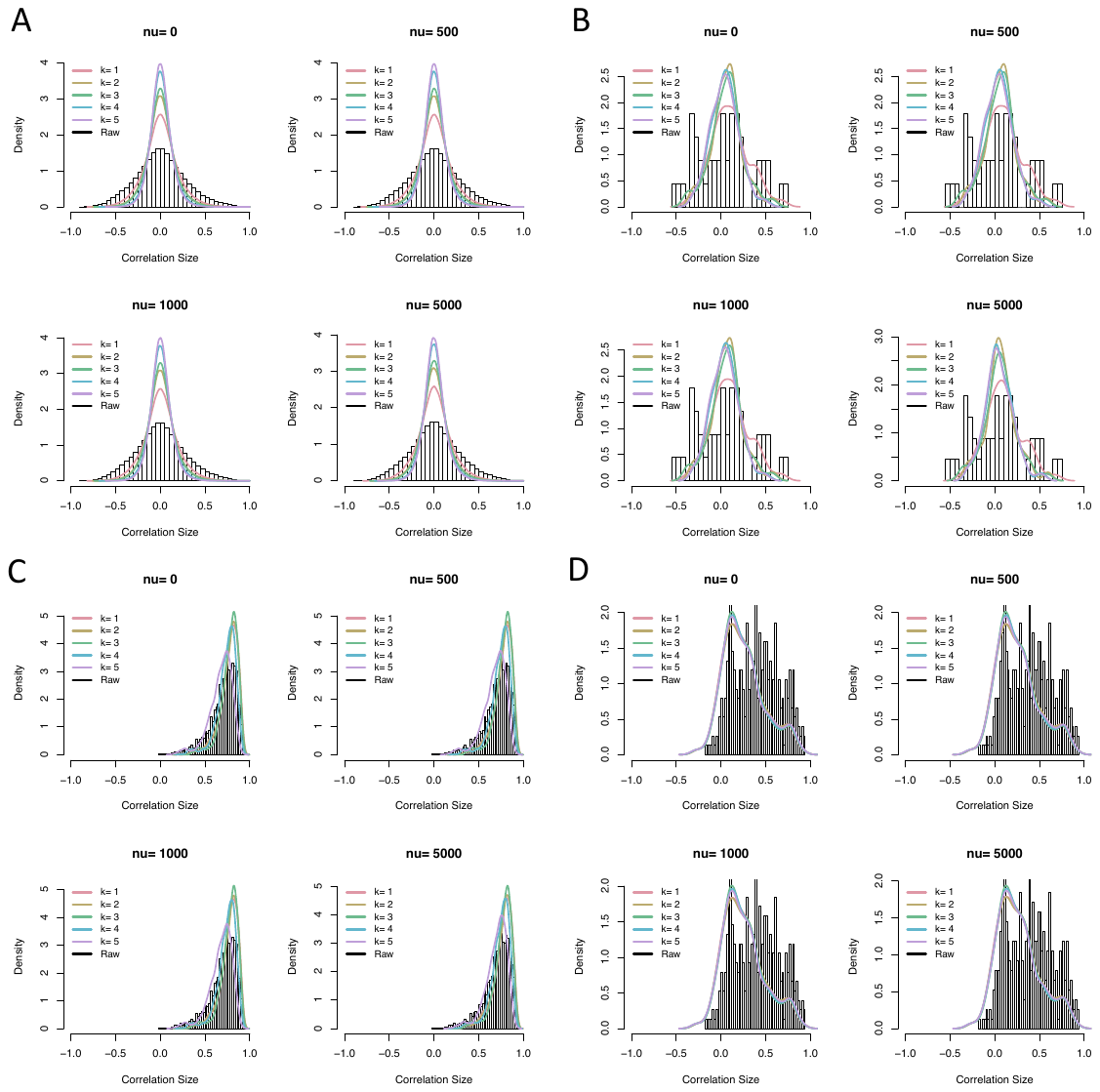

**Figure S10**. Distribution of gene-gene correlations at different RUVCorr parameter settings. Tissue = heart-left ventricle. The background histogram shows the distribution of gene-gene correlations for each gene set, calculated using the unadjusted data. A) Random genes, B) Sodium channel genes, C) Ribosome genes, D) Major histocompatibility complex (MHC) genes

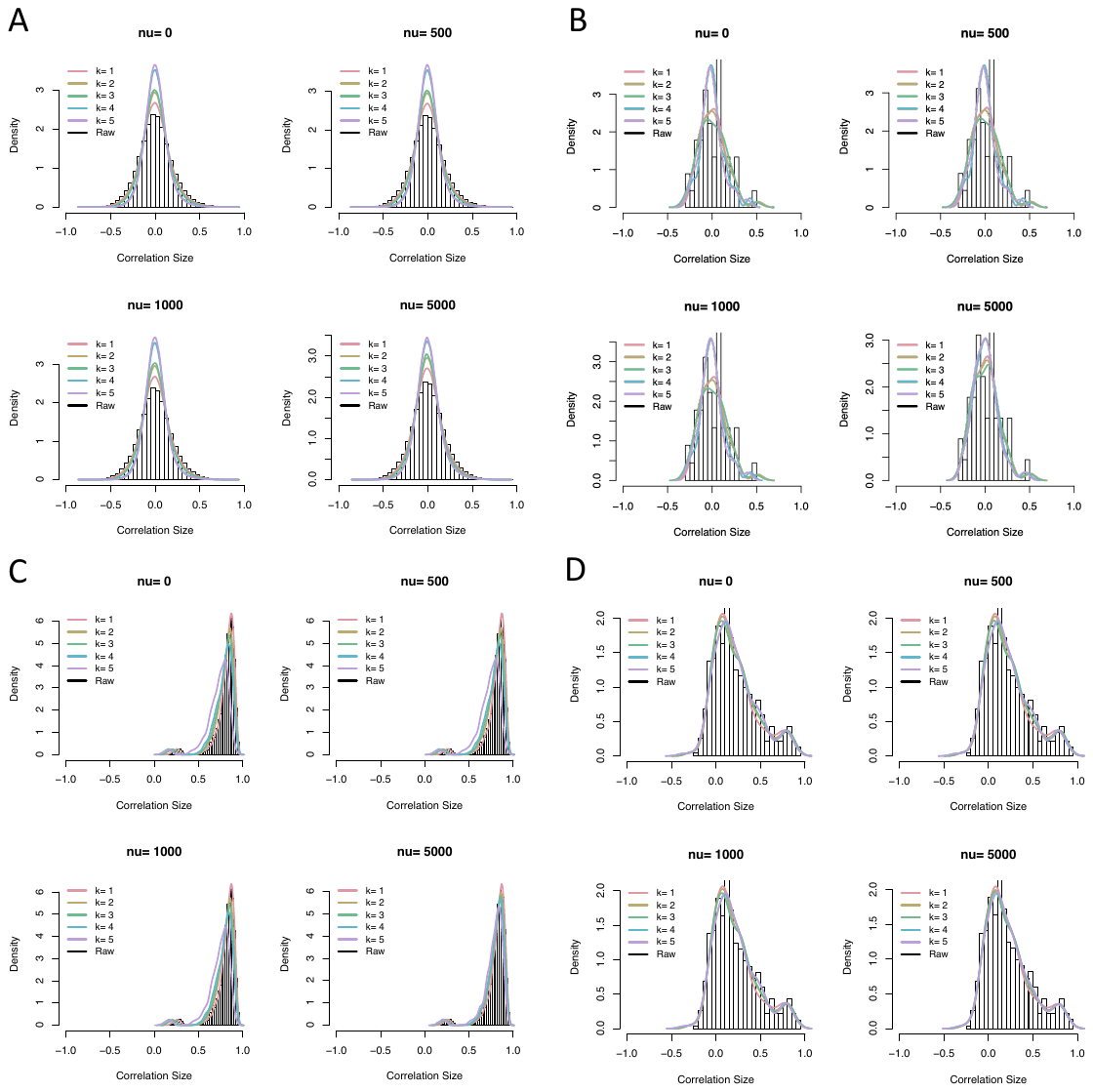

**Figure S11**. Distribution of gene-gene correlations at different RUVCorr parameter settings. Tissue = spleen. The background histogram shows the distribution of gene-gene correlations for each gene set, calculated using the unadjusted data. A) Random genes, B) Sodium channel genes, C) Ribosome genes, D) Major histocompatibility complex (MHC) genes

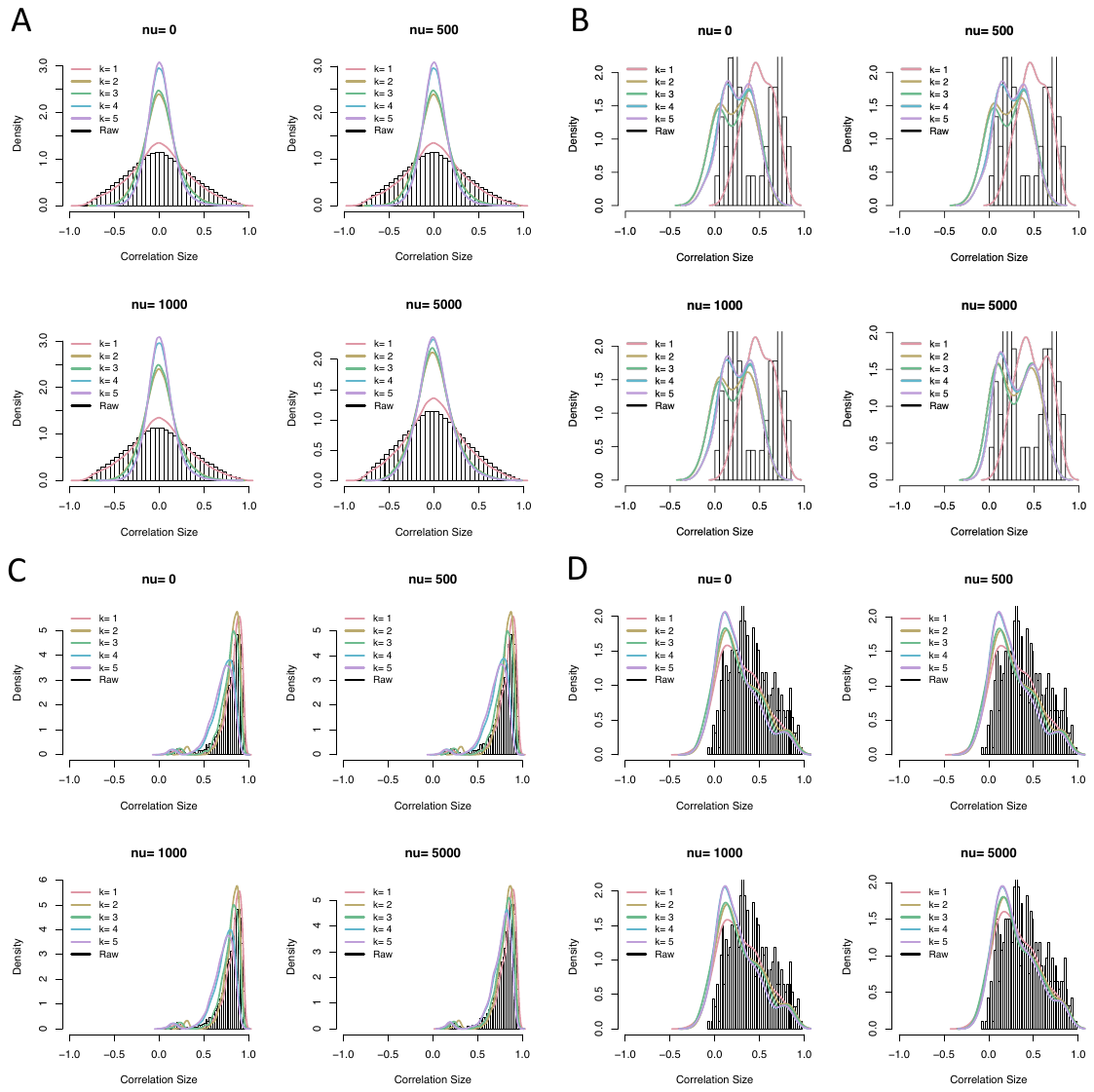

**Figure S12**. Distribution of gene-gene correlations at different RUVCorr parameter settings. Tissue = small intestine. The background histogram shows the distribution of gene-gene correlations for each gene set, calculated using the unadjusted data. A) Random genes, B) Sodium channel genes, C) Ribosome genes, D) Major histocompatibility complex (MHC) genes

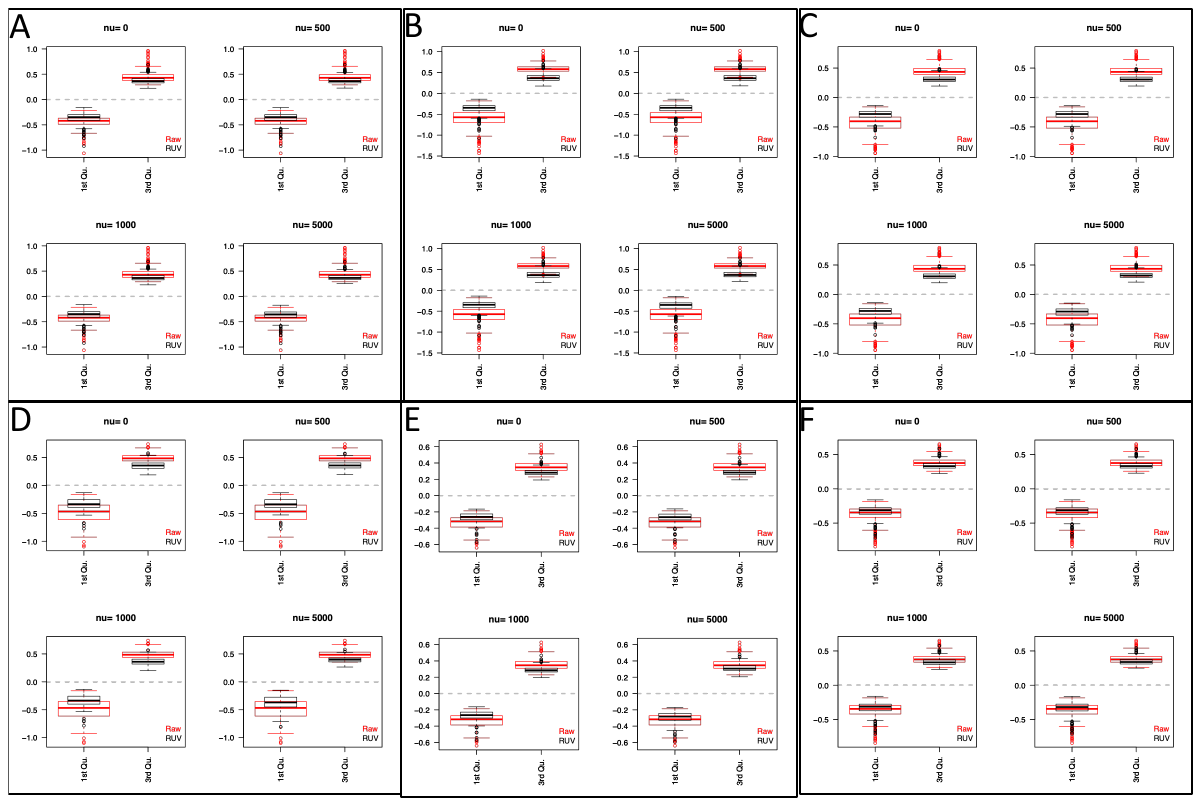

**Figure S13**. Histogram plots of relative log expression at optimal k parameter and various nu parameters. Each figure includes boxplots of the first and third quartile of difference between gene expression and the median for raw and RUV-adjusted data. A) Skeletal muscle (k=2), B) Whole blood (k=4), C) Heart-left ventricle (k=4), D) Small intestine (k=2), E) Spleen (k=4), F) Adipose (k=3).

**Table S1**. **Sample characteristics, data correction parameters, and AUC score gene pairs**

| Tissue | Number of Samples | Number of Genes | RUVCorr | | CONFETI | PC | AUC Score | |
| --- | --- | --- | --- | --- | --- | --- | --- | --- |
|  |  |  | k | nu | Number of IC's used as confounding factors | Number of PCs included in adjustment | True positive pairs | True negative pairs |
| Whole Blood | 670 | 20358 | 4 | 1000 | 348 / 454 | 49 | 15113 | 8010 |
| Adipose | 581 | 23976 | 3 | 500 | 386 / 457 | 49 | 18139 | 9987 |
| Skeletal muscle | 706 | 21084 | 2 | 500 | 453 / 533 | 53 | 17128 | 5092 |
| Small intestine | 174 | 26247 | 2 | 1000 | 105 / 126 | 12 | 17230 | 11316 |
| Spleen | 227 | 25537 | 4 | 1000 | 161 / 191 | 28 | 17345 | 9992 |
| Heart-left ventricle | 386 | 21412 | 4 | 1000 | 246 / 292 | 32 | 16793 | 5413 |

**Table S2. Covariates included in known covariate adjustment, per tissue.**

| **Tissue** | **Covariates** |
| --- | --- |
| Whole blood | Sex, age, PC1-5, RNA integrity number, total number of transcripts with at least 5 exon mapping reads, total ischemic time, chimeric pairs, exonic rate, fragment length stdev, intergenic rate, number of reads aligning to rRNA regions, intronic rate, mapped unique rate of total, alternative alignments, number of split reads |
| Adipose | Sex, age, PC1-5, autolysis score, RNA integrity number, total ischemic time, total number of genes with at least 5 exon mapping reads, fragment length stdev, number of all reads aligning to ribosomal RNA regions, failed vendor QC check, intronic rate, mapped unique rate of total, alternative alignments, base mismatch rate, number of End 1 reads sequenced in the sense direction, rRNA rate |
| Skeletal muscle | Sex, age, PC1-5, RNA integrity number, total ischemic time, End 2 mapping rate, chimeric pairs, intragenic rate, exonic rate, number of reads aligning to rRNA regions, total number of transcripts with at least 5 exon mapping reads, alternative alignments, mean fragment length, base mismatch rate, rRNA rate |
| Small intestine | Sex, age, PC1-5, autolysis score, RNA integrity number, total ischemic time, time spent in PAXgene fixative, chimeric pairs, intragenic rate, total number of genes with at least 5 exon mapping reads, End 1 mismatch rate, number of reads aligning to rRNA regions, failed vendor QC check, intronic rate, mapped unique rate of total, alternative alignments, mean fragment length, base mismatch rate, number of End 1 reads sequenced in sense direction, rRNA rate, End 1 mapping rate, End 2 % sense |
| Heart-left ventricle | Sex, age, PC1-5, RNA integrity number, total ischemic time, intragenic rate, exonic rate, failed vendor QC check, total number of transcripts with at least 5 exon mapping reads, alternative alignments, mean fragment length, split reads, base mismatch rate, number of End 1 reads sequenced in the sense direction, End 1 % sense, rRNA rate, End 1 mapping rate, End 2 % sense |
| Spleen | Sex, age, PC1-5, RNA integrity number, autolysis score, total ischemic time, time spent in PAXgene fixative, End 2 mapping rate, chimeric pairs, End 1 mismatch rate, intergenic rate, number of all reads aligning to rRNA regions, failed vendor QC check, total number of transcripts with at least 5 exon mapping reads, intronic rate, End 2 antisense, alternative alignments, base mismatch rate, End 1% sense, rRNA rate, End 1 mapping rate, End 2% sense |
